# Effective use of genetically-encoded optical biosensors for profiling signalling signatures in iPSC-CMs derived from idiopathic dilated cardiomyopathy patients

**DOI:** 10.1101/2022.09.06.506800

**Authors:** Kyla Bourque, Ida Derish, Cara Hawey, Jace Jones-Tabah, Kashif Khan, Karima Alim, Alyson Jiang, Hooman Sadighian, Jeremy Zwaig, Natalie Gendron, Renzo Cecere, Nadia Giannetti, Terence E. Hébert

## Abstract

Dilated cardiomyopathy (DCM) is a cardiovascular condition that develops when the left ventricle of the heart enlarges, compromising its function and diminishing its capacity to pump oxygenated blood throughout the body. After patients are diagnosed with DCM, disease progression can lead to heart failure and the need for a heart transplantation. DCM is a complex disease where underlying causes can be idiopathic, genetic, or environmental. An incomplete molecular understanding of disease progression poses challenges for drug discovery efforts as effective therapeutics strategies remain elusive. Decades of research using primary cells or animal models have increased our understanding of DCM but has been hampered due to the inaccessibility of human cardiomyocytes, to model cardiac disease, *in vitro*, in a dish. Here, our goal is to leverage patient-derived hiPSC-CMs and to combine them with biosensors to understand how cellular signalling is altered in DCM. With high sensitivity and versatility, optical biosensors represent the ideal tools to dissect the molecular determinants of cardiovascular disease, in an unbiased manner and in real-time at the level of single cells. By characterizing the pathobiology of dilated cardiomyopathy in a patient-specific manner using high content biosensor-based assays, we aim to uncover personalized mechanisms for the occurrence and development of DCM and as a pathway to development of personalized therapeutics.

## Introduction

Cardiomyopathy is a heart muscle disease characterized by increased myocardial stress caused by the diminished capacity of the heart to adequately pump oxygenated blood throughout the body. There are five types of cardiomyopathies: dilated (DCM), hypertrophic (HCM), restrictive (RCM), arrhythmogenic right ventricular dysplasia (ARVD) and unclassified which includes Takotsubo cardiomyopathy. Each disease impacts patient prognosis differently. Irrespective of the etiological form: genetic or non-genetic, cardiomyopathy patients progress towards heart failure and many eventually require heart transplantation. Unfortunately, 50% of individuals diagnosed with heart failure (HF) succumb to this disease within 5 years of diagnosis. Of interest, dilated cardiomyopathies are characterized by left- or bi-ventricular dilation/enlargement and systolic dysfunction. A large fraction of DCM patients have a genetic etiology with 25% of these patients harbouring mutations in the titin gene. Besides the giant sarcomeric filament titin, mutations in several other sarcomere-related genes result in DCM (reviewed in (1,2)). DCM pathology can also be caused due to infection, autoimmunity as well as chemical and toxin exposure (3), such as alcohol or heavy metals (4,5). Further, some prescription (6) or illicit drugs (7) have been associated with DCM. Of significance, certain chemotherapeutic agents have important side effects that progress towards a DCM phenotype either acutely or years after administration (8–11). In addition, endocrine dysfunction and certain metabolic syndromes have been linked with the disease (12,13). Dilated cardiomyopathy can occur secondarily to neuromuscular causes (14). However, many patients are diagnosed with idiopathic forms, in part, due to inadequate genetic testing (15). Thus, with such heterologous underlying etiologies, challenges arise in developing novel therapies to treat DCM or cardiomyopathies in general-necessitating a more personalized attention.

A better understanding of the underlying molecular determinants that drive disease progression or therapeutic response is needed. The advent of induced pluripotent stem cells (iPSCs) allows the conversion of somatic cells, including patient-derived samples, into stem cells that can later be differentiated into functional cardiomyocytes (CMs) or other relevant cell types for phenotypic studies (16,17). Numerous reports have demonstrated the power of human-derived iPSC-CMs to model various aspects of DCM, particularly genetic and chemotherapy-induced cases (**Table 1**). Several reviews have also been published showing how iPSC-CMs have been used to model genetic or sarcomeric cardiomyopathies (18–22).

**Table 1.**
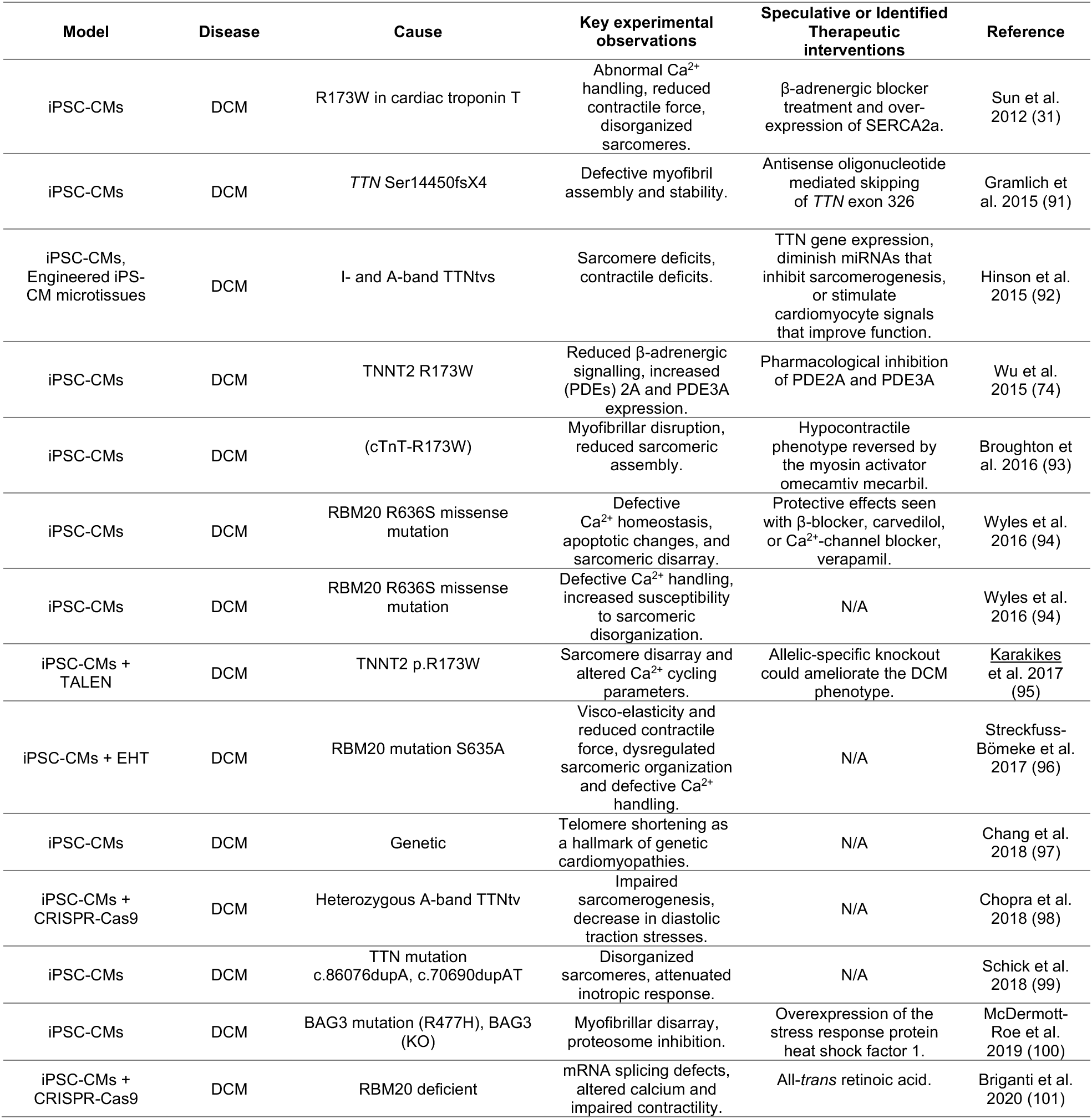
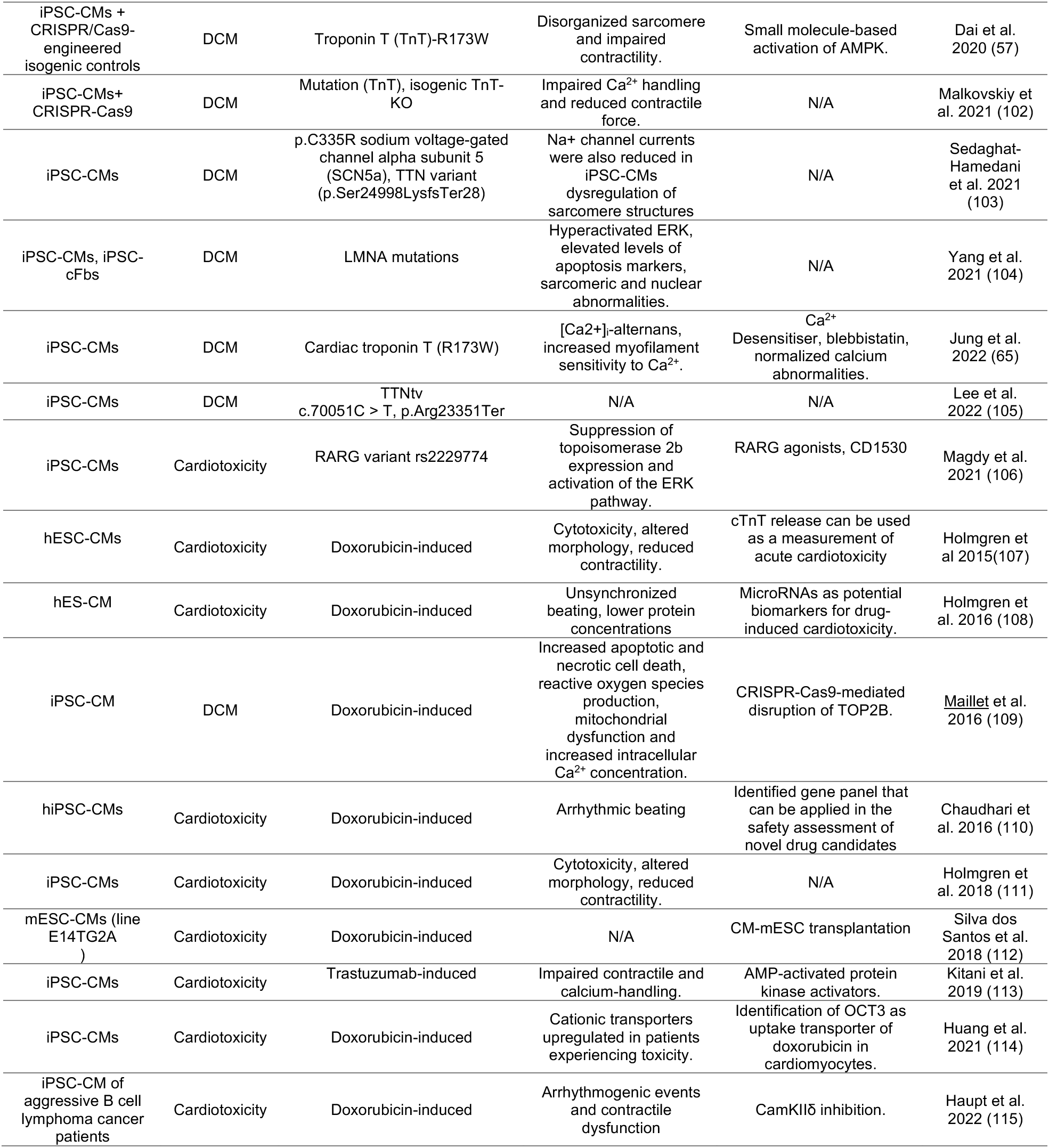
Non-exhaustive literature search of iPSC-CM based *in vitro* models of dilated cardiomyopathy.

To begin to understand the specific molecular events that drive idiopathic DCM disease progression, we have established an iPSC-based program for personalized cardiovascular medicine enabled by access to patient-derived blood samples followed by iPSC generation. Our biobank of iPSCs when differentiated into cardiomyocytes are combined with an optical biosensor-based screening platform to identify patient specific signalosomes as means to better track idiopathic disease mechanisms. Genetically-encoded optical biosensors (GEBs) are chromophore-based and can be used to probe cellular responses to specific stimulations (23,24).

To obtain a mechanistic understanding of DCM at the molecular level, we have established proof of concept data that use biosensors to model idiopathic DCM, *in vitro*, in a dish. The biosensors used here target pathways downstream G protein-coupled receptors (GPCRs) as the activation of these signalling cascades is interconnected with the pathobiology and pharmacotherapy of dilated cardiomyopathy. We explored protein kinase A (PKA) (25,26) and extracellular signal-regulated protein kinase (ERK_1/2_) activity (27) as well as calcium handling in patient and control iPSC-derived cardiomyocytes (iPSC-CMs) as an approach to uncover the unknown molecular anonymities causing idiopathic DCM. PKA and calcium handling were selected as important regulators of cardiac contractility (28) while ERK_1/2_ signalling is an important cell survival signal in the myocardium (29,30). We are specifically interested in assembling detailed patient-specific cellular signalling profiles to obtain a first glance at the diversity or similarities across the entire patient and control cohort. Ultimately, we would like to uncover whether features of the disease process are shared amongst male and female subjects or whether age, etiology, etc. are determining factors of the molecular profiles being gathered. At the outset, we want to know whether the idiopathic nature of the disease will *seep* into the iPSC-CM model as well as genetic DCM which has been relatively well recapitulated *in a dish* (31). With the diverse etiologies of our patient cohort, described below, we will have the inherent capacity to examine the molecular fingerprint of idiopathic, genetic, familial and chemotherapy induced DCM. Our novel approach can also assess whether iPSC-CM maturity influences *in vitro* modeling of idiopathic disease. Based on the current logistical model, maturation does not seem to influence DCM disease modeling when assessing genetic etiologies (31). However, some conflicting evidence has been reported such as when performing clinical trials in a dish (32,33). Yet, to what extent will this be the case for idiopathic DCM? Since biosensors allow for repeated measures, signal profiles can be assessed at distinct timepoints (day 30, 45, 60) to control for cardiomyocyte maturity while removing potential batch effects by assaying the same cells. This data will feed the development of personalized interventions through improved understanding of individual disease mechanisms and therapeutic responses. Below, we describe our patient population and show preliminary data that supports using optical biosensors for idiopathic disease modelling.

## Materials and Methods

### Recruitment of patients and control subjects

Male and female patients as well as control subjects were recruited at the cardiology clinic at the McGill University Health Center. Control subjects were either healthy family members of the patients or participants with no known cardio-pulmonary disease. All enrolled participants provided informed consent to be a part of the ‘Heart-in-a-Dish” project with REB approval HID-B/2020-6362. Each individual completed a questionnaire that recorded information concerning their biological sex, age, ethnicity, cardiovascular family history, smoking, cholesterol, past cancer diagnosis, current medications and a genesis praxy gender questionnaire.

### Generation and validation of hiPSCs from healthy control volunteers and DCM patients

#### Reprogramming of PBMCs into induced pluripotent stem cells

Peripheral blood mononuclear cells (PBMCs) were grown in StemSpan SFEMII supplemented media (STEMCELL Technologies, CA) which are distinct since they can be observed to float in culture. Once PBMCs were grown to sufficient numbers, at least 1×10^6^ PBMCs, the Epi5™ Episomal iPSC Reprogramming Kit (Thermofisher, USA) was used along with the Neon Transfection System (Invitrogen) to introduce Oct4, Sox2, Lin28, Klf4, and L-Myc into the cells using the following parameters: 3 pulses, 10 ms, 1650 V. Once electroporated, cells were seeded onto freshly prepared Matrigel-coated plates (Corning), to help with the attachment of the iPSCs. Reprogramming, as per (34), was regarded as successful when floating cells began to attach to the cell culture dish, observed on approximately day 7 after electroporation. Once attached, iPSC colonies slowly expanded and were mechanically split under the microscope. Briefly, a 20- or 22-gauge needle was used to delineate the border of the colony, and a grid was then made for colony splitting. iPSC colonies were selected after gross morphological inspection under a brightfield microscope. High quality iPSCs that were picked displayed large nuclei, were tightly packed, and colony borders had a clear delineation, without any differentiation or fibroblast-like cells at the borders (16,35).

#### Validation of pluripotency and quality of iPS cell lines

All iPSC lines will be subsequently validated (36–38) by 1) colony morphology; 2) alkaline phosphatase; 3) qRT-PCR and immunocytochemistry assays of pluripotency markers (e.g., Nanog, SSEA4, SOX2, Klf4, Tra 1-60). iPSC colonies at passages 6 and onward were immunostained to assess expression of core pluripotency genes at the protein level. For immunofluorescence studies, hiPSCs were replated onto Matrigel-coated 24-well plates, and colonies were fixed with 4% PFA once they measured approximately at 100 – 500 μm in diameter. Next, colonies were permeabilized using 0.5% Triton X in PBS for 15 minutes at RT. Cells were then stained with primary antibodies overnight, at 4°C, for nuclear pluripotency markers OCT4 (Cell Signaling, #2750) and NANOG (Cell Signaling, #4903), as well as surface markers SSEA-4 (Cell Signalling, #4755) and TRA-1-60 (Cell Signaling, #4746). The following day, iPSC colonies were co-stained with secondary antibody AlexaFluor-488, anti-rabbit or anti-mouse, for 1 hour at RT. Representative images were captured using a fluorescent microscope. If the localization and intensity of the staining of each respective antibody was found to be appropriate across all colonies imaged, this confirmed pluripotency.

Further, to functionally validate the ability of iPSC to differentiate into the three germ layers-ectoderm, mesoderm, and endoderm, a trilineage differentiation assay was performed (R&D Systems, SC027B). The expression of Otx2 was assessed as it is a nuclear protein responsible for the primitive streak stage of embryonic development – it is closely related and expressed in the ectoderm, which gives rise to neurons, eyes, skin, etc. In addition, the expression of Brachyury was examined as it is a transcription/mesodermal factor expressed in tissues fated to become the heart, bone marrow, etc. Brachyury is mostly found in the nucleus. Lastly, Sox17, found in the nucleus and cytoplasm, was studied as it is an important factor during endoderm formation throughout the gastrulation process. Based on the manufacturer’s instructions, iPSC colonies were seeded into 2×3 wells of a 24-well plate, and grown according to the R&D Systems’ protocol. Each well (in duplicate) was differentiated and stained with the respective markers above. This assay essentially allowed for the qualitative assessment of the quality and potential of our iPSC lines to differentiate.

To ensure chromosomal integrity, hiPSC lines were screened for normal karyotype by G-banding technique. Briefly, iPSC colonies were seeded into two Matrigel coated 60-mm tissue culture dishes and harvested at 50-60% confluency. The morning of the harvest, Colcemid (ThermoFisher, 15212012) was added onto the iPSCs at two different concentrations; 0.10 μg/mL and 0.15 μg/mL and incubated at 37°C for 30 min. Cells were then washed three times with PBS prior to being incubated with Accutase solution. Dissociated iPSCs were collected and spun down at 1000 RPM for 10 min in a 15 mL conical tube. If the cell pellet was between 2 to 4 mm in size, the protocol was continued, otherwise pellets were discarded. Once the supernatant was removed, leaving approximately 0.5 mL, a pre-warmed, 37°C, 0.075 M KCl hypotonic solution was added onto the pellet and the tube was inverted ~7 times. In the fume hood, cells were prefixed by adding 8 drops of RT Carnoy’s fixing solution (3 methanol: 1 glacial acetic acid) to the KCl solution. The tube was then inverted 10 times followed by centrifugation at 1000 RPM for 10 min. In the fume hood, the supernatant was aspirated until 0.5 mL and the pellet was resuspended by flicking the tube. Cold fixing solution, 1 mL, was then added and the tube was inverted 10 times. Cells were then incubated at −20°C for 1 hour. After being inverted a couple times, cells were spun down at 1000 RPM for 20 min at 4°C. The supernatant was removed until 0.5 mL and the pellet was resuspended using flicking motion before being resuspended in 14 mL cold Carnoy’s fixing solution. Cells were then shipped to Cytogenomics Facility at the Hospital for Sick Kids in Toronto, Ontario, Canada for karyotyping. All iPSC lines were also screened for mycoplasma contamination using a mycoplasma detection kit (ThermoFisher, 4460623). Once the test came back negative, iPSC lines were established within our biorepository and subsequently differentiated into cardiomyocytes.

#### Differentiation of hiPSCs into functional cardiomyocytes

Control and patient-derived iPSCs were differentiated into cardiomyocytes following the previously published GiWi protocol with minor modifications (39). To achieve a 95% confluent monolayer of iPSCs, 500,000 cells were seeded onto Matrigel (Corning, 354277) coated 24-well dishes in mTeSRPlus media supplemented with 10 μM Rho kinase inhibitor, Y-27632-HCl (Selleckchem, S1049). The following day, spent media was removed and 1 mL fresh mTeSRPlus media was added onto the iPSC and cells were returned to the incubator. On day 0 of the GiWi protocol, Wnt signalling was activated by supplementing the RPMI 1640 + B27 minus insulin media (ThermoFisher A1895602) with 12 μM Wnt activator or GSK-3 inhibitor, CHIR99021 (Cayman Chemical, 13122). The time was recorded to ensure the media would be exchanged 24 hours later with fresh RPMI 1640 supplemented with B27 minus insulin. On day 3, Wnt signalling was inhibited using the IWP2 compound (Selleckchem S7085). Precisely 48 hours later, on day 5, the spent media was removed and exchanged with RPMI 1640 supplemented with B27 minus insulin. On day 7 and day 10 of the protocol, the media was changed for RPMI 1640 supplemented with regular B27 (ThermoFisher, 17504001). At this time, spontaneous beating iPSC-CMs should be observed under a brightfield microscope using 10X or 20X magnification. Starting at day 10-12, metabolic selection was performed and the iPSC-CMs were starved from glucose for 4-6 days. RPMI 1640 without glucose, supplemented with B27 as well as with 4 mM lactate was used to purify the cardiomyocyte population (40). Wells embedded with monolayers of spontaneously beating cardiomyocytes were replated into fibronectin-coated 6-well dishes and maintained in RPMI 1640 media supplemented with B27 until day 35-40 where 30,000 iPSC-CMs were re-seeded into optical bottom, black 96-well plates for cellular signalling experiments (Nunc, 165305).

#### Transduction of hiPSC-CMs with adeno-associated viruses

Adeno-associated viruses (AAVs) used in this study were produced by the Neurophotonics Platform Viral Vector Core at Laval University, Québec. To minimize freeze-thaw cycles, AAVs were aliquoted in low-retention Eppendorf tubes and stored for long-term at −80°C. Post-thawing, AAVs were kept at 4°C for a maximum of 7-10 days. To determine the optimal serotype to transduce our iPSC-CM cultures, we used the serotype selection kit offered by the Neurophotonics Platform (data not shown). The kit contains AAV serotypes 1, 2, 5, 6, 8, 9, Rh10, DJ, DJ8, php.B, php.eB, php.S and retro for testing. Based on these results, iPSC-CMs were transduced with biosensors packaged within AAV serotype 6, which had previously been reported to be effective at infecting cardiomyocytes (41,42). During transduction, AAVs were kept on ice before diluting them in RPMI 1640 + B27 media. To minimize excessive cell debris carried over during cell seeding into the microwell dish, cardiomyocyte cultures were washed three times with basal RPMI 1640 media before being transduced. A multiplicity of infection (MOI) of 5000, representative of 5000 viral genomes per cell was used for all experiments. Post virus addition, iPSC-CM cultures were returned to the 37°C incubator and kept there for a minimum of 72 hours to allow for sufficient biosensor expression.

#### Single-fluorophore intensity-based and dual-colour FRET imaging

On the day of the experiment, cardiomyocyte cultures were visualized under a phasecontrast microscope to confirm that the iPSC-CMs were of good quality. Essentially, we confirmed that the iPSC-CMs were spontaneously beating and appeared to have a cardiomyocyte-*like* morphology. If the cells were deemed healthy, the media was exchanged for pre-warmed, 37°C, assay buffer, HBSS without phenol red, with calcium, magnesium, and sodium bicarbonate (Wisent 311-513-CL). Cells were washed three times prior to leaving the cells in 90 μL HBSS for the assay. The microwell plate was then returned to a humidified atmosphere of 37°C with 5% CO2 for 45 minutes to an hour before imaging. This permitted the iPSC-CMs sufficient time to re-equilibrate in the assay buffer post-washing. Meanwhile, the temperature control settings of Perkin Elmer’s Opera PHENIX high-content screening system were turned on. The microscope would thus have sufficient time to warm up to 37°C with 3% CO_2_ for live-cell imaging. Concurrently, drugs were prepared at the benchtop at a 10-fold more concentration as 10 μL would be added in each well, which is equivalent to a 10-fold dilution. For EKAR-NLS based experiments, FRET imaging was collected using a 20X air objective using a 425 nm laser for excitation of the CFP donor. Emissions were detected with filters at 435-515 nm (CFP-donor) and 500-550 nm (YFP-acceptor). For the ExRai-AKAR2-NLS biosensor, images were acquired using a 20X air objective using a 375 nm and 480 nm laser for excitation and 500-550 nm emission filter. In the experimental setup within Perkin Elmer’s Harmony software, parameters were set, where the automated microscope would capture an initial basal, ligand independent, measurement. Subsequently, the assay microplate was ejected from the system, and drug stimulations were conducted manually. To minimize cellular perturbations, we were cautious not to move/displace the microplate during agonist addition. The microscope then continued collecting 7 measurements post-drug stimulation for another 70 minutes at 10-minute intervals as programmed.

#### Single-fluorophore intensity-based and dual-colour FRET analysis

Post-imaging raw image files were imported into Perkin Elmer’s Columbus image analysis software. As biosensors were localized to the nucleus, iPSC-CM nuclei were identified using the ‘find nuclei’ feature. Subsequently, nuclear morphology features, including nuclei roundness and area were calculated. The morphology criteria permitted the sorting of the nuclei’s to then select a population of ‘healthy’ nuclei without including unwanted auto-fluorescent cell debris. A roundness and size threshold were set to remove cellular debris which had rough edges and were smaller in size compared to a typical cell’s nucleus. To calculate the CFP/YFP FRET ratio for EKAR-NLS biosensor, the intensity properties of the donor and acceptor fluorophores were computed. The same calculations were performed by Columbus to calculate the GFP excitation ratio for the ExRai-AKAR2-NLS biosensor. Data of each individual nucleus that was categorized as ‘healthy’ was then exported as text files for further analysis in R, as the following.

Single nuclei datasets were treated in R based on previously published single-cell analytical approach with minor modifications (43). Briefly, cardiomyocyte nuclei that appeared in all 8 timepoints (t=0 and post-stimulation) were carried forward. As nuclei tend to drift slightly once the microplate is loaded back into the microscope post drug addition, a threshold of fluorescence intensity and distance was set for each nucleus. To be considered as the same object (nucleus), the fluorescence intensity could deviate by ≤20%. Nuclei that did not conform to this parameter were excluded from the analysis. To compute the change in FRET in response to drug addition, a ΔFRET parameter was calculated. ΔFRET represents an individual’s nuclei FRET relative to basal, ligand independent FRET. This was then converted into a percentage change in FRET (%ΔF/F). In this percentage change in FRET, the denominator (F) was computed by averaging the basal FRET across all nuclei in the same microwell. The TSclust package (44) in R was then used to cluster the cells by the magnitude of response over time. In this package, the ‘pam’ function was used. For the single nuclei clustering, patient/control or male/female nuclei were merged into a single dataset. This allowed for consistency within the patterns and clusters identified within the two populations that were being assessed. Performing the clustering algorithm on separate dataset may result in distinct clusters being identified. This method would assure the clusters identified would be matched in the two datasets. Following the clustering, the merged patient/control or male/female dataset were split and further plotted as heatmaps for visualization using pheatmap.

#### Calcium imaging and analysis

Single cell calcium handling was measured using RGECO-TnT, a red-shifted single-colour intensiometric biosensor that localizes to troponin T in the myofilament (45). iPSC-CMs were seeded at a density of 20,000 cells per microwell of a black, optical bottom, 96-well plate. Approximately three days later, iPSC-CMs were washed three times with basal RPMI 1640 media before being transduced with AAV2/6-RGECO-TnT using an MOI of 5000. Prior to recording calcium transients, iPSC-CM cultures were inspected under a phase-contrast microscope to ensure cells were healthy. Then, media was exchanged with RPMI 1640 without phenol red supplemented with B27 to minimize background caused by autofluorescent phenol red. Spontaneous, ligand independent calcium transients in iPSC-CMs were captured at 10.4 frames per second for 15 seconds using a Zeiss Axio Observer fully automated inverted microscope with a Zeiss 20× PLAN APOCHROMAT (NA 0.8), X-Cyte 120 LED light source, and FS-14 RFP filter set (560/26 nm excitation, 620/60 nm emission, 565 dichroic mirror). For the duration of the experiment, hiPSC-CMs were kept in a temperature- and gas-controlled chamber, mimicking physiological conditions, 37°C and 5% CO_2_. The open-source image processing software, Image J, was used to trace single cells from image stacks. Analysis of recorded videos was performed using using a custom Transient Analysis app in OriginPro 2021b v9.8.5 (OriginLab). This software has the built-in capacity to determine features of calcium transients including but not limited to the calcium transient frequency, time to peak, peak amplitude, time between peaks, transient duration, area under the curve, time to reach 50% baseline, time to reach 90% baseline and upstroke velocity. Analysis was performed on a single-cell basis. Statistical analysis was done using GraphPad Prism v9.3.1 with a Welch’s ANOVA test and Dunnett’s multiple comparisons controlling for unequal variances.

## Results and Discussion

### Establishment of control and DCM iPSC lines

To develop a more personalized approach to cardiac care, we created a biorepository of patient samples from those diagnosed with dilated cardiomyopathy as well as age-matched healthy control subjects (**Fig. 1 & Suppl. Table 1, 2**). Enrolled patients had different suspected primary etiologies of their respective dilated cardiomyopathic phenotypes ranging from idiopathic, viral, genetic, chemotherapy-induced etc. providing us with a comprehensive patient population to sample from (**Fig. 1A**). The large majority of our patients also have a left ventricular ejection fraction of less than 40%, a feature characteristic of systolic dysfunction (**Fig. 1B**). Thus far, we have recruited 228 individuals who consented to be part of the “Heart-in-a-Dish” project. The current patient population includes men and women of different ages and ethnicities as well as healthy individuals that will serve as control subjects (**Fig. 1C-F**). Based on available patient data, the family history of cardiac disease is higher in the patient pool compared to control (**Suppl. Table 1**). Risk factors associated with HF are also more prevalent in the DCM patient population. Most patients have been prescribed pharmacological agents that target neurohormones; the β-adrenergic or renin-angiotensin-aldosterone system (RAAS) such as β-blockers, ACE inhibitors, or a combination of angiotensin receptor and a neprilysin inhibitor (**Suppl. Table 2**).

**Figure 1.**
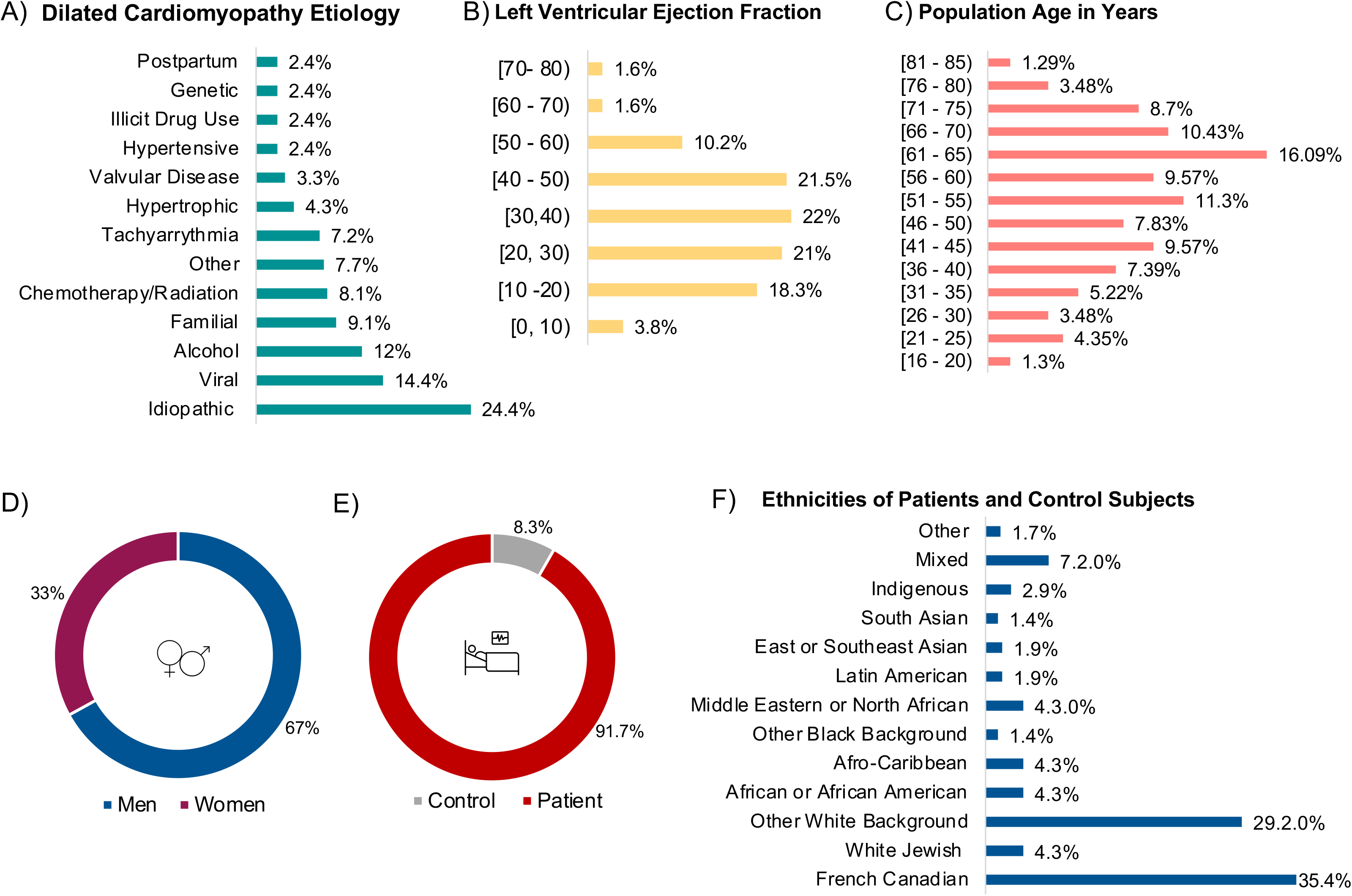
Overview of general characteristics of the patient and control population enrolled in the “Heart-in-a-Dish” study. **(A) Bar chart illustrating the percentage per primary suspected etiologies of the registered patient cohort.** Heterologous DCM etiologies range from idiopathic, familial, genetic, chemotherapy, alcohol, viral among others. **(B) Graph demonstrating the percent of patients with different left ventricular ejection fractions (LVEF).** A large fraction of patients fall within a LVEF percentage of less than 40%, a typical feature of systolic dysfunction. **(C) Age and (D) Biological sex of patient and control population.** Patients of all ages have been recruited in the study as well as a well-balanced ratio of males and females. **(E) Patients as well as control subjects have been recruited as part of the study.** An approximate 10:1 ratio has been achieved when enrolling patient and control subjects respectively. **(F) Various ethnicities of the patient and control subjects.** Thirteen ethnicities have been self-proclaimed by the study participants when they filled in the questionnaire during enrollment.

To model dilated cardiomyopathy in a dish, patient blood was extracted using vein-puncture and stored in the biobank for subsequent analysis. CD34^+^ cells were isolated from the buffy coat and reprogrammed into iPSCs using episomal plasmids or non-integrating Sendai-virus. Successful reprogramming into a primordial pluripotent state was confirmed by immunofluorescence for pluripotency markers, as well as tri-lineage differentiation (**Fig. 2A-C**). Once the “stemness” of the iPSC lines was validated followed by normal karyotype analyses (**Suppl Fig. 1**), hiPSCs were differentiated into cardiomyocytes using a defined protocol that modulates Wnt signalling (**Fig. 3A**) (46). Spontaneous contracting monolayers were observed for all patient and control lines that entered the cardiac differentiation program. The cardiomyocyte lineage was further validated by gene and protein expression studies (**Fig. 3B, C**). Troponin I3 expression increased with time in culture indicating maturation of the iPSC-CMs, and staining with anti-a-actinin2 showed that most of the cells were in fact cardiomyocytes. The expression of the β-adrenergic receptors was increased over time in culture, demonstrating the functionality and ability of iPSC-CMs to respond to external stimuli (**Fig. 3D**). Once cardiomyocyte identity was confirmed and after 4 weeks in culture, we virally introduced optical biosensors using adeno-associated viruses (**Fig. 3E**) as tools to shed light on underlying DCM disease mechanisms, as described below.

**Figure 2.**
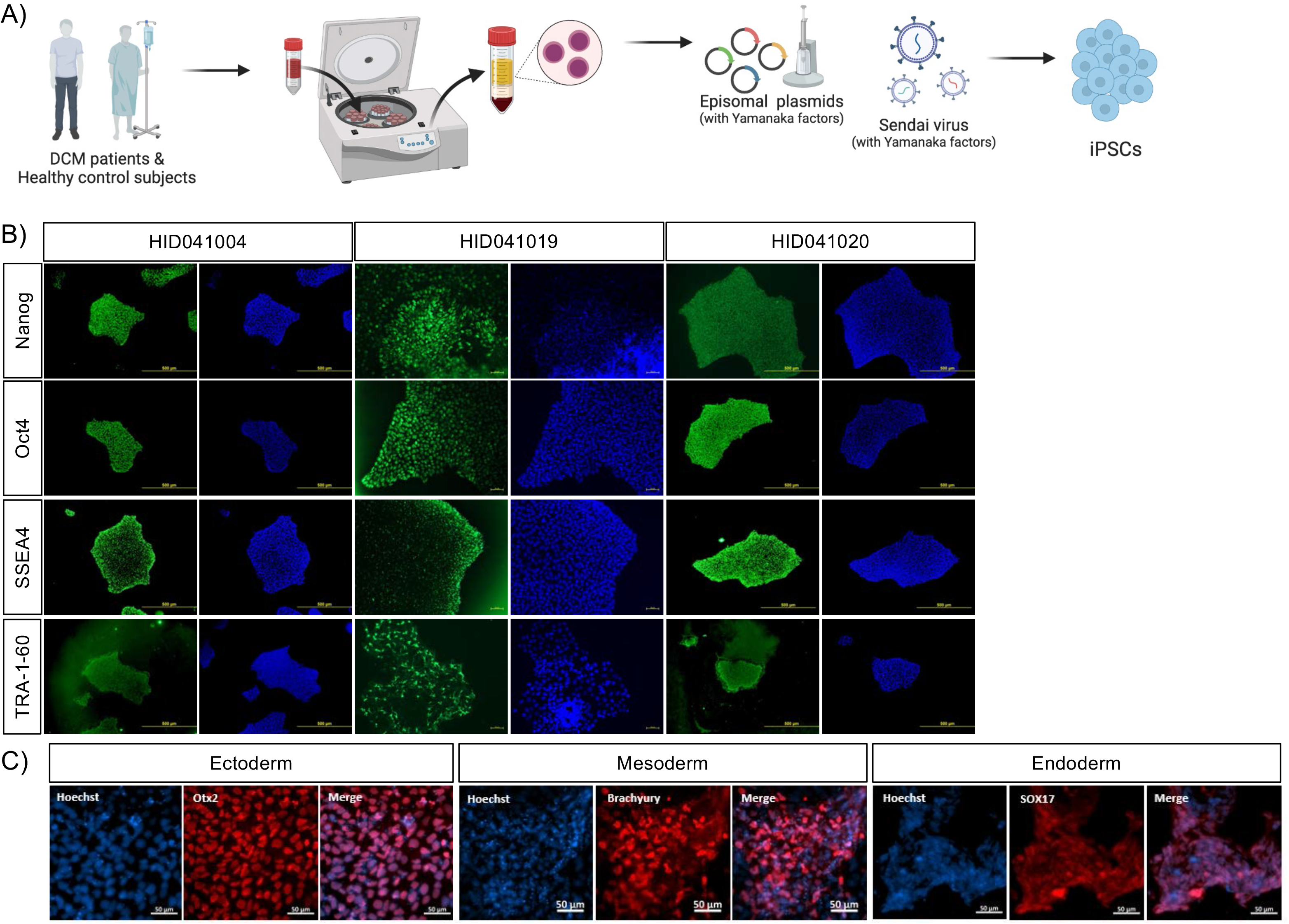
Reprogramming and validation of patient- and control derived iPSCs. **(A) Schematic illustrating the process required to reprogram PBMCs into iPSCs using either episomal plasmids or non-integrating Sendai virus techniques.** The process of reprograming PBMCs into iPSCs takes on average 2-3 months. To some extent, the efficiency of reprogramming was inversely proportional with age, with samples sourced from older individuals exhibiting lower reprogramming efficiency, requiring at times, repeated attempts. **(B) Immunofluorescence images depicting protein expression of key pluripotency markers: Nanog, Oct4, SSEA4 and TRA-1-60.** As depicted, OCT4 and NANOG co-localize with Hoechst stains and SSEA-4/TRA-1-60 are found at the cell surface. **(C) Tri-lineage differentiation of a representative iPSC cell line.** Fluorescent microscopy images demonstrate the pluripotential of the iPSC line to differentiate into the three germs layers, ectoderm (Otx2), mesoderm (Brachyury) and endoderm (Sox17).

**Figure 3.**
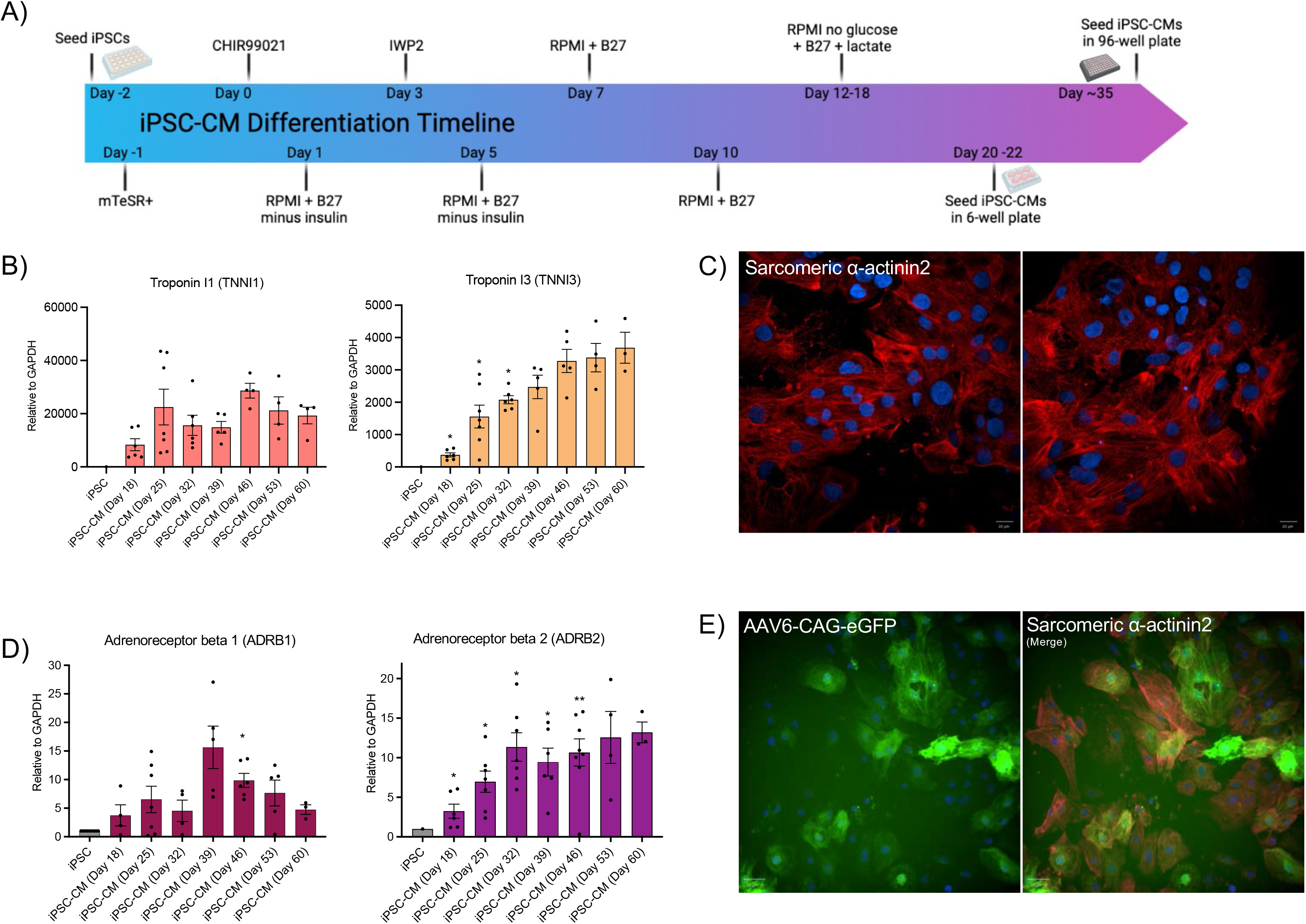
Differentiation of patient and control iPSCs into functional cardiomyocytes. **A) Schematic illustration of the differentiation protocol to generate cardiomyocytes from iPSCs.** Through the temporal modulation of Wnt signalling, the protocol yields spontaneously contracting cardiomyocytes within 7-10 days. **(B) RT-qPCR gene expression analysis revealed cardiomyocyte fate of differentiated iPSCs.** Expression of troponin isoforms relative to housekeeping gene GAPDH of three or more independent differentiations over time. **(C) Immunofluorescence images demonstrating protein expression of a sarcomere-related gene.** iPSC-CMs stained with α-actinin2 followed by secondary antibody Alexa-647 and Hoechst nuclear dye validates cardiomyocyte identity. **(D) RT-qPCR gene expression analysis demonstrating that iPSC-CMs are signalling competent.** RT-qPCR gene expression of β_1_- and β2-adrenergic receptors establishes that cardiomyocytes can transduce signals through cell surface receptors that are also critical targets for DCM pharmacotherapy. **(E) iPSC-derived cardiomyocytes are effectively transduced using AAV serotype 6.** Fluorescent microscopy images depicting the delivery of eGFP into iPSC-CMs using AAV2/6. The green colour of eGFP is a well matched when iPSC-CMs are co-stained with sarcomeric marker α-actinin2. Correspondingly, all biosensors in this study were packaged in serotype 6. Mean ± SEM, * p<0.5, \*\**p* < 0.01.

### Biosensors for cardiovascular disease modelling

Over the past decade, genetically-encoded biosensor (GEB) technology has undergone significant development. Biosensors have been designed to probe for the activity of more than 100 different signalling pathways, many of which are cardiac relevant (47,48) (**Fig. 4A**). GEBs typically fall into two broad categories, either being based on resonance energy transfer (RET) or on the intensity of biosensor emission (reviewed in (49,50)). Briefly, RET is a quantum optical process that relies on the non-radiative transfer of energy from a donor to an acceptor. Compatibility between donor and acceptor probes requires the donor’s emission spectrum to overlap with the excitation wavelength of the acceptor. This energy transfer event is dependent on distance (up to a maximum of 10 nm) as well as orientation (reviewed in (51–54)). However, certain versions are less dependent on orientation (27). Addedly, RET biosensors can either have a bioluminescent donor (BRET) or a fluorophore as donor (FRET). BRET biosensors typically suffer from weaker signals but have higher signal to noise ratios compared to FRET equivalents (55). FRET however is more amenable to single cell imaging and microscopy applications. In contrast, intensiometric biosensors report on analyte concentrations or effector activation in accordance with fluctuations in the magnitude of fluorescence intensity emitted by a single fluorophore. In this context, the fluorescence of intensiometric biosensors is quenched in the absence/presence of the molecule being probed and changes to the degree of pathway activation (increase/decrease) alters the intensity of the fluorescence signal emitted. Since intensiometric sensors monitor signalling by recording the emission of a single wavelength, their main advantage is their ability to be multiplexed.

**Figure 4.**
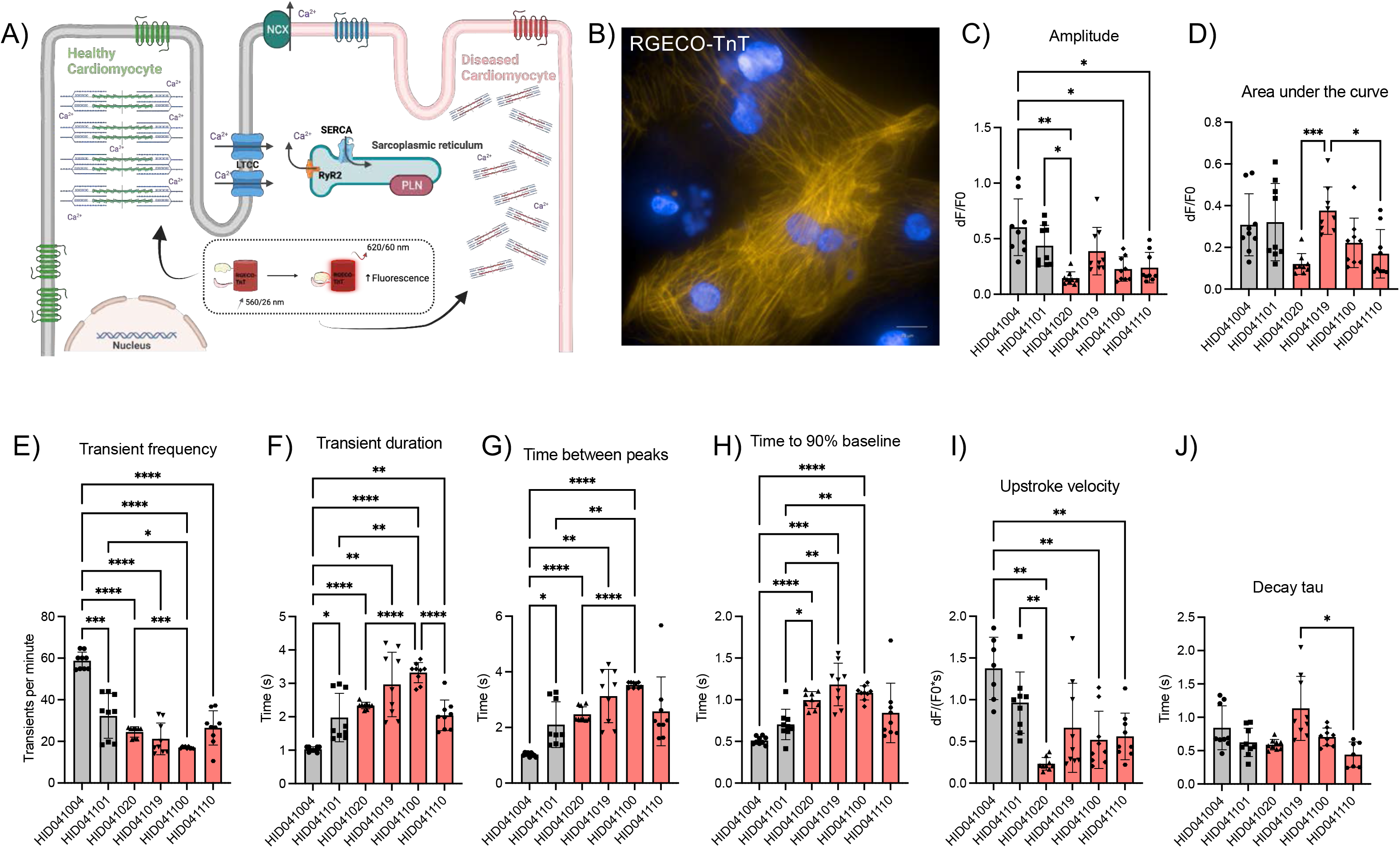
Calcium imaging as measured in iPSC-CMs derived from healthy control subjects and idiopathic DCM patients. **(A) Calcium handling in the heart.** Since calcium is a critical regulator of cardiomyocyte contractility, abnormal calcium cycling has been observed to result in impaired cardiac functions as seen in the failing heart and dilated cardiomyopathy. (**B) Representative fluorescent microscopy image of the red-shifted RGECO-TnT biosensor in iPSC-cardiomyocytes.** The myofilament localization of RGECO-TnT is clearly observed with the yellow/orange fluorescence. Several calcium transient properties were measured including **(C)** calcium transient amplitude, **(D)** area under the curve, **(E)**transient frequency, **(F)** transient duration, **(G)**time between peaks, **(H)** time to 90% baseline, **(I)** upstroke velocity, and **(J)** decay tau in both control subjects (gray: HID041004 and HID041101) and idiopathic DCM patients (red: HID041020, HID041019, HID041100, HID041110), n= 9 cells from 1-2 biological replicates. * = p<0.05, ** = p<0.01, *** = p<0.001, **** = p<0.0001.

Biosensors are excellent tools for the detection of numerous biological interactions in realtime and in live cells. When used in human iPSCs and their differentiated derivatives, as well as in patient-derived samples, biosensor methods have the potential to explore disease causative mechanisms. For example, genetically encoded biosensors have facilitated the study of calcium and kinase signalling in iPSC-CMs derived from hypertrophic cardiomyopathy (45) and dilated cardiomyopathy (56,57) patients respectively. GEBs have also helped resolve electrophysiological changes in stem cell derived CMs while revealing their higher throughput capabilities when using GEBs compared to traditional methods (58,59). However, several studies still rely on indicator dyes such as Fluo-3 and Fluo-4 AM to study calcium handling in patient derived iPSC-CMs (31,60). Even if the use of indicator dyes does present with certain advantages, such as ease of use without the need of transfection, biosensor technology represents a more versatile approach. Most notably, the catalog of GEBs allows us to dissect changes in the disease- and patientspecific signalosomes investigating the activity of ion channels, protein kinases as well as a long list of protein effectors. Importantly, GEBs tend to result in less cellular toxicity compared to the use of dyes and can be more easily targeted to intracellular compartments (discussed in (58,59,61,62)). Lastly, GEBs are amenable to repeated measures allowing to track cardiomyocyte function in long-term measurements and in response to chronic stressors, a feature that is useful as cardiovascular disease evolves overtime. To our knowledge, biosensor-based studies have only been used in diseased iPSC-CMs with inherited or strong genetic etiologies.

### Combining optical biosensors with iPSC-CMs

Much evidence points to dysregulated signalling pathways in the development of dilated cardiomyopathy. Considering that DCM shows a phenotypic spectrum with inherited and environmental etiologies, the molecular underpinnings of the disease may be more complex than previously anticipated. To study DCM pathogenesis in iPSC-CM monolayers, we choose genetically-encoded biosensors that interrogate ERK_1/2_ and PKA signalling as well as calcium handling as these have been shown to be altered in DCM. We also elected to follow changes in signalling at the single-cell level. Recently single cell approaches have gained prominence as they are being performed at larger scale and tend to generate richer datasets. Bulk analyses may allow for high-throughput biosensor assays and are generally less computationally intensive compared to single-cell analytical approaches. Yet, single cell approaches provide in-depth analysis of the complex behaviours of all cells within the overall population. Individual cells process information differently as their intracellular and extracellular milieus are distinctive, including neighboring cells and growth factor concentrations. Consequently, no cell perceives information the same way and the cellular state has also been reported to impact cellular signalling behaviors of single cells (63).

We hypothesized that by examining heterogenous cell populations, at the single cell level, different cell behaviors and sub-populations would emerge when assessing their functionality using biosensor technologies. Single cell analyses allow to better understand the variability that shapes the population as a whole. In line with this, we applied two single cell analytical pipeline to dissect biosensor data collected using the Opera Phenix automated high content microscope from Perkin Elmer or the Zeiss Axio Observer (43). The Opera Phenix was used to track ERK_1/2_ and PKA signalling overtime while the faster imaging speed of the Zeiss Axio Observer was used to collect calcium handling data. For the ERK_1/2_ and PKA assays, Columbus image analyses software then allowed us to examine the images extracting details relating to each individual cell or nuclei. Integrated within analytical software such as Columbus, it becomes possible to find individual nuclei or cells using the donor channel. Further, it is also possible to extract morphological features of individual cells/nuclei. This morphological information can be used in the next steps to correlate signalling with cell- or nuclei-specific morphologies (43). For example, nuclei size and roundness can be computed in addition to cell size, roundness, length, width, and length-to-width ratio. Extracting this information allows users to deduce whether a correlation exists between these morphological features and the signalling signature of each individual cell (43). Once these features have been extracted from the collected images, text files can be imported into R for further processing. Here, we used R as it is an open-source free software environment that allows processing of large multivariate datasets. For calcium imaging, raw images were imported into Image J for feature extraction prior to being analysed in Origin Lab software. Below, we explain how our analytical approach previously used on primary neuronal cultures (43) as well as single-cell calcium imaging can help assess signalling in iPSC-CMs derived from control or idiopathic DCM patients.

### Exploring disease-specific cellular vulnerabilities by probing cardiomyocyte calcium handling

Numerous reports have described impaired calcium handling and corresponding contractile function in DCM. For example, reduced calcium transients were observed in human ventricular muscle strips sourced from DCM hearts compared to non-failing heart samples (64). In a study using iPSC-CMs derived from a patient carrying a R173W mutation in Troponin T, CMs displayed delayed times-to-peak Ca^2+^ transients, showing calcium release from the SR was compromised as well as delayed reuptake kinetics (65). TnT-R173W iPSC-CMs have also been shown to have reduced calcium transient amplitudes speculated because of reduced SR Ca^2+^ storage (66). In addition to dysregulated calcium handling and contractility, sarcomere misalignment is another common feature observed in DCM cardiomyocytes (66). In line with this, shorter sarcomere lengths as well as reduced contractile amplitude have been reported in TnT-R173W iPSC-CMs (67). These data suggest a destabilized calcium-troponin network in DCM. Overall, numerous reports have identified impaired calcium handling as a phenotypic abnormality in iPSC-based models of DCM.

Calcium handling deficits have been linked with numerous mutations associated with the development of hypertrophic and dilated cardiomyopathy with mutations being found in cardiomyocyte cytoskeletal and contractile proteins (**Table 1.0**) (68). The role of calcium in the heart is extensive as the heart contracts and relaxes via excitation-contraction, E-C coupling, and involves tight modulation of intracellular calcium concentrations (**Fig. 4A**). To discern molecular level differences between DCM patients as well as in comparison to healthy controls, the assessment of cardiomyocyte calcium handling becomes a significant phenotypic endpoint. To accomplish this, [Ca^2+^]_i_ was measured at the myofilament using a red-shifted intensiometric genetically-encoded calcium indicator called RGECO-TnT (**Fig. 4B**) (45). Our proof-of-concept single-cell results indicate that differences in myofilament-localized calcium handling can be captured between DCM patients and control subjects as well as between DCM patients. Features of calcium transients such as time between peaks, transient frequency, and upstroke velocity displayed a general trend where iPSC-CMs from DCM patients exhibited slower spontaneous calcium transients than iPSC-CMs derived from healthy controls (**Fig. 4C-J**). Furthermore, calcium transients from DCM patients were shown to have lower amplitudes and longer transient durations than healthy controls. Features such as time to 90% baseline, transient duration, and area under the curve exemplified that this biosensor, RGECO-TnT, when expressed in iPSC-CMs can be used to detect differences in myofilament-localized calcium handling between DCM patients, some of which were classified as idiopathic (**Fig. 4C-J**). After confirming these results in a larger patient sample, it will be possible to test whether the abnormal calcium handling features observed here can be rescued using pharmacological agents. Overall, calcium is a useful proxy that can be used to assess cellular vulnerabilities and how DCM pathogenesis affects individual cardiomyocytes on a patient-specific basis.

### Exploring disease-specific signalling signatures in iPSC-CMs through the measurement of PKA activation profiles

Protein kinase A (PKA) is an important effector in regulating cardiomyocyte function and behavior as it is involved in regulating contractility, growth, and metabolism. For example, in failing human hearts, PKA-mediated phosphorylation of troponin has been reported to be reduced compared to healthy hearts (69). A similar observation was reported in *Gα_q_* transgenic mice suggesting PKA has a role in the development of DCM (70). Cardiac apoptosis driven by norepinephrine has also been shown to be facilitated by a PKA-dependent pathway (71). Transgenic mice expressing the catalytic subunit of PKA have also been observed to develop dilated cardiomyopathy (72). The hyperphosphorylation of PKA substrates was also shown to be detrimental to heart function (72). Inhibiting PKA phosphorylation of RyR2 effectively reversed the DCM phenotype and abnormal calcium handling in mdx mice (73). Interestingly, in iPSC-CMs, the PKA response to isoproterenol was shown to be attenuated in a genetic DCM line compared to control (74). Similarly, iPSC-CMs carrying a TnT-R173W mutation were shown to bind PKA at reduced levels compared to wildtype troponin supporting the reduced contractility observed in DCM (57). Further, cAMP levels at sarcomere myofilaments were shown to be increased, indicative of a compensatory mechanism within cardiomyocytes to increase PKA responsiveness (57). Based on this accumulated evidence, pointing towards a role of PKA in DCM progression, we sought to examine whether PKA activation profiles were informative of disease by comparing data gathered from a control male subject and a male patient (**Fig. 5**).

**Figure 5.**
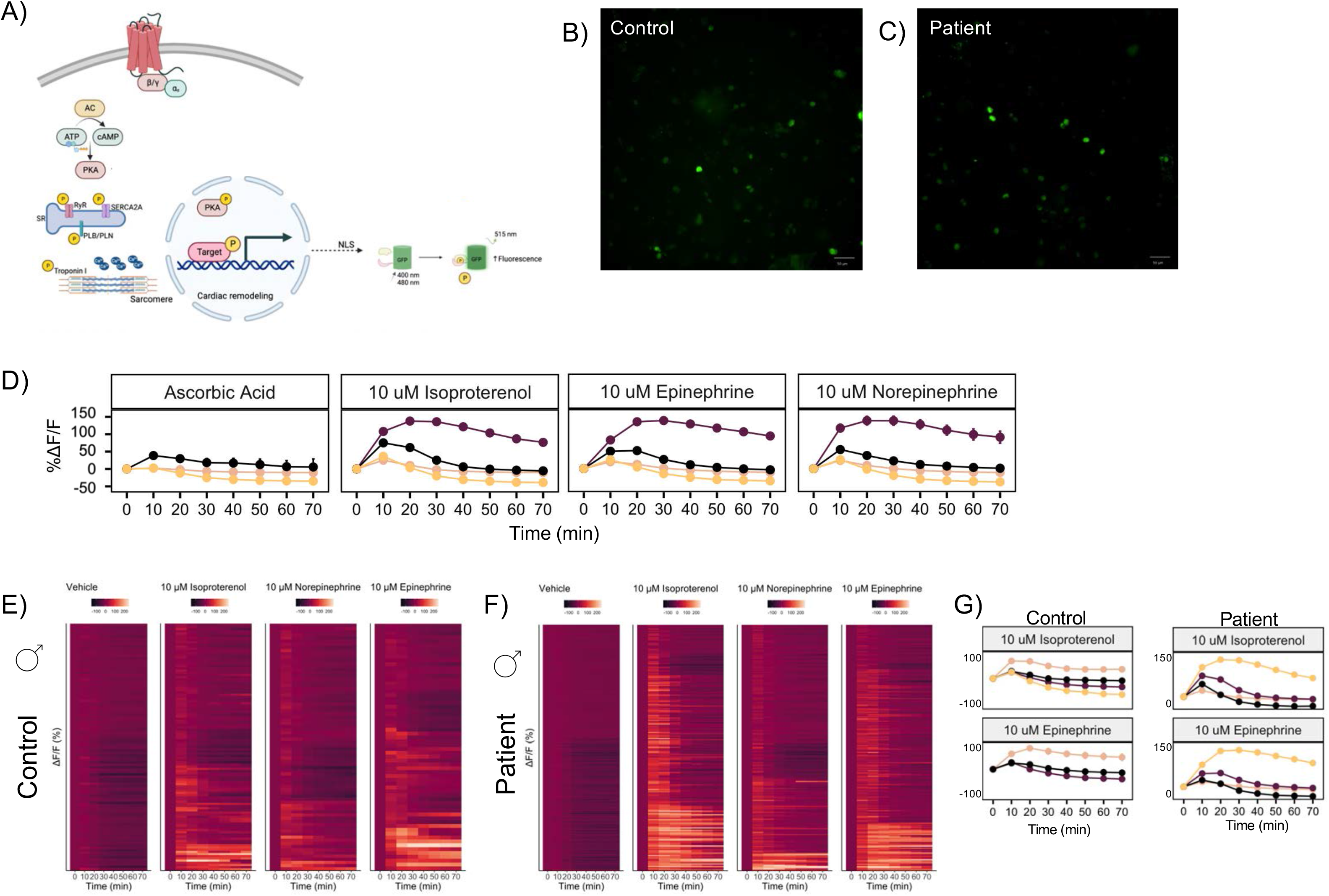
Biosensors to illuminate disease-specific signalling. **(A) Schematic illustration of genetically-encoded biosensor to interrogate nuclear protein kinase A (PKA) activity.** PKA is activated downstream adrenergic receptors and regulates numerous aspects of cardiac physiology and pathophysiology including the function of sarcomeric proteins, cardiac contractility, and adverse remodelling. **(B) Fluorescent microscopy images of ExRai-AKAR2-NLS.** Representative images display nuclear localization of the biosensor in **(B)** patient and **(C)** control iPSC-CM nuclei. **(D) Summary of distinct clusters identified in response to adrenergic agonists.** iPSC-CMs were observed to fall within four different clusters depending on their behaviors. Nuclei were found to exhibit sustained (plum), or transient responses (black) while other nuclei failed to produce a measurable response (light pink) or exhibited a decrease in activity from baseline (yellow). **(E-F) Representative heatmap representing single cell results of intensity-based biosensor ExRai-AKAR2-NLS in (E) control and (F) patient iPSC-CMs.** The signalosome of two male subjects, control and patient were compared post stimulation by either vehicle, isoproterenol, epinephrine, or norepinephrine. Images were collected pre- and post-drug stimulation. Data was collected for 70 minutes post-drug addition. The male patient demonstrated a ‘hyper-activated’ PKA phenotype as a larger proportion of cells responded to the tested agonists. **(G) Response clusters for control and patient iPSC-CMs.** When plotted independently, patient clusters are indicative of a ‘hyperresponsive’ phenotype with a more robust increase in %ΔF/F compared to control (see y-axis).

To reveal whether PKA signalling was perturbed in DCM patients, we transduced both patient and control iPSC-CMs using AAV2/6-ExRai-AKAR2-NLS, an intensity-based, single (green) colour biosensor that reports of PKA activity in the nucleus (25) (**Fig. 5A-C**). This biosensor has a large dynamic range and its brightness allows for robust responses to be measured. After the recording of a basal measurement, iPSC-CMs were stimulated with saturating doses of agonists that target β-adrenergic (isoproterenol, epinephrine, norepinephrine) and α-adrenergic receptors (norepinephrine, epinephrine). Agonist-induced changes in PKA activity were subsequently recorded. All nuclei were then run through a clustering algorithm to uncover patterns within the data, that cannot be detected with an untrained eye (**Fig. 5D**). Four clusters were noted, demonstrating four distinct behaviours. Nuclei that exhibited a sustained response to agonists as well as transient responses were observed. Some nuclei did not respond to the drugs tested while other nuclei exhibited a decreases overtime in response compared to baseline. This clustering feature within our analysis demonstrates the ability to identify novel sub-populations of cells that exhibit distinct patterns for further analysis (44). To visualize agonist induced changes in PKA activity, single nuclei responses were plotted as a heat map (**Fig. 5E, F**). In the control subject, approximately 40% of cells exhibited a sustained or transient response to adrenergic drugs. In the patient nuclei, the ratios of these clusters were evidently distinct in response to isoproterenol (**Fig. 5F**). The two largest clusters observed appeared to represent nuclei populations that exhibited a sustained or transient increase in PKA activity over time. We also observed a distinct pattern in response to norepinephrine, demonstrating that different β-agonists result in different single nuclei signalling signatures. In comparison to the control, the data seems to suggest that the male patient’s iPSC-CMs displays a ‘hyper-active’ phenotype as almost 70% of iPSC-CM nuclei led to a robust measurable response compared to control. Similar observations were observed in response to epinephrine, albeit to a lesser degree.

Here, we demonstrated the capacity of biosensors to uncover distinct PKA activation profiles in a DCM patient. As DCM and HF are associated with impaired calcium handling, this hyperactive phenotype can be speculated to present due to signalling cross-talk. Data collected from this patient demonstrates reduced basal calcium transients and calcium transient amplitudes (**Figure 4**). As cAMP and calcium signals exhibit a certain level of cross-talk, lower basal calcium may also impact the basal cAMP-PKA axis causing the PKA response to β-agonists to appear as hyperactivation. With lower basal cAMP-PKA, there may be a larger *‘range’* for PKA activity to increase. When sustained, a hyper-adrenergic phenotype can result in increased phosphorylation of PKA substrates resulting in cardiomyocyte remodelling.

### Exploring the effect of biological sex on ERK_1/2_ activity in DCM iPSC-CMs

Mitogen-associated protein kinases (MAPKs) are important regulators of molecular events in many cells. Myocardial activation of ERK_1/2_ is complicated as its activation can result in cellular survival signals as well as maladaptive hypertrophic growth and fibrosis. The nature, kinetics and tone of the signal are important to consider as well as their localization and intensity. MAPK signalling has been identified to be abnormal in cardiomyopathies and HF. For instance, ERK_1/2_ has been shown to be abnormally elevated in sections of explanted human DCM hearts measured using a phospho-specific antibody (75). An ERK_1/2_ inhibitor, selumetinib has been demonstrated to prevent cardiac ERK_1/2_ activation, which led to positive outcomes on cardiac function in mice (75). The I61Q cTnC DCM mutation was correlated with cytosolic ERK_1/2_ activity and elongated cardiomyocytes while a HCM mutation were associated with nuclear translocation of ERK_1/2_ and demonstrated features of concentric hypertrophy (76). Hyperactivated p38, ERK_1/2_ and JNK signalling has been detected in models of LMNA-dependent dilated cardiomyopathy (77,78). Mitogen activated protein kinase inhibitors have been demonstrated to improve the LMNA dilated cardiomyopathic phenotype (79–82). However, ERK_1/2_ signalling was not altered in guinea pig left ventricular cardiomyocytes modeling DCM mutations of the thin-filament regulatory proteins (83). Thus, there is a clear phenotypic spectrum in terms of signalling when assessing different mutations and models of dilated cardiomyopathy.

To assess ERK_1/2_ activity in iPSC-CMs, we transduced our cultures with AAV2/6-EKAREV-NLS (27) (**Fig. 6A**). As shown for the male patient, the FRET biosensor was well expressed, expressing donor and acceptor fluorophores at comparable intensities (**Fig. 6B, C)**. Upon β-agonist stimulation, a transient ERK_1/2_ activity profile appeared in the male patient compared to the iPSC-CMs derived from a female patient (**Fig. 6D, E**). This female patient presented with a larger cluster of nuclei which exhibited a decrease in ERK_1/2_ activity post stimulation of adrenergic drugs compared to baseline. With a prominent basal ERK_1/2_ signature in females, it can be speculated that females could be more protected from cell death pathways compared to males (84). Yet, ERK_1/2_ has also been associated with adverse cardiac remodelling and may be suggestive of distinct basal disease manifestations. A quantitative analysis of the FRET signals, through the generation of calibration curves, may help further elucidate the differences in basal ERK_1/2_ activity. ET-1 treatment led to the most robust increases in ERK_1/2_ activation compared to all other drugs tested, in both sexes. A secondary assessment of cell size would help illuminate whether a hypertrophic gene program is underway ensuing an eccentric hypertrophic phenotype. The male patient also displayed a small cluster of iPSC-CMs experiencing a transient ERK_1/2_ activation in response to Ang II. We also tested SI and SII, two compounds known for their β-arrestin biased nature. The latter representing ligand properties of interest due to their believed cardioprotective effects (85–87). Yet, in the male patient, these drugs exhibited a similar response profile compared to Ang II.

**Figure 6.**
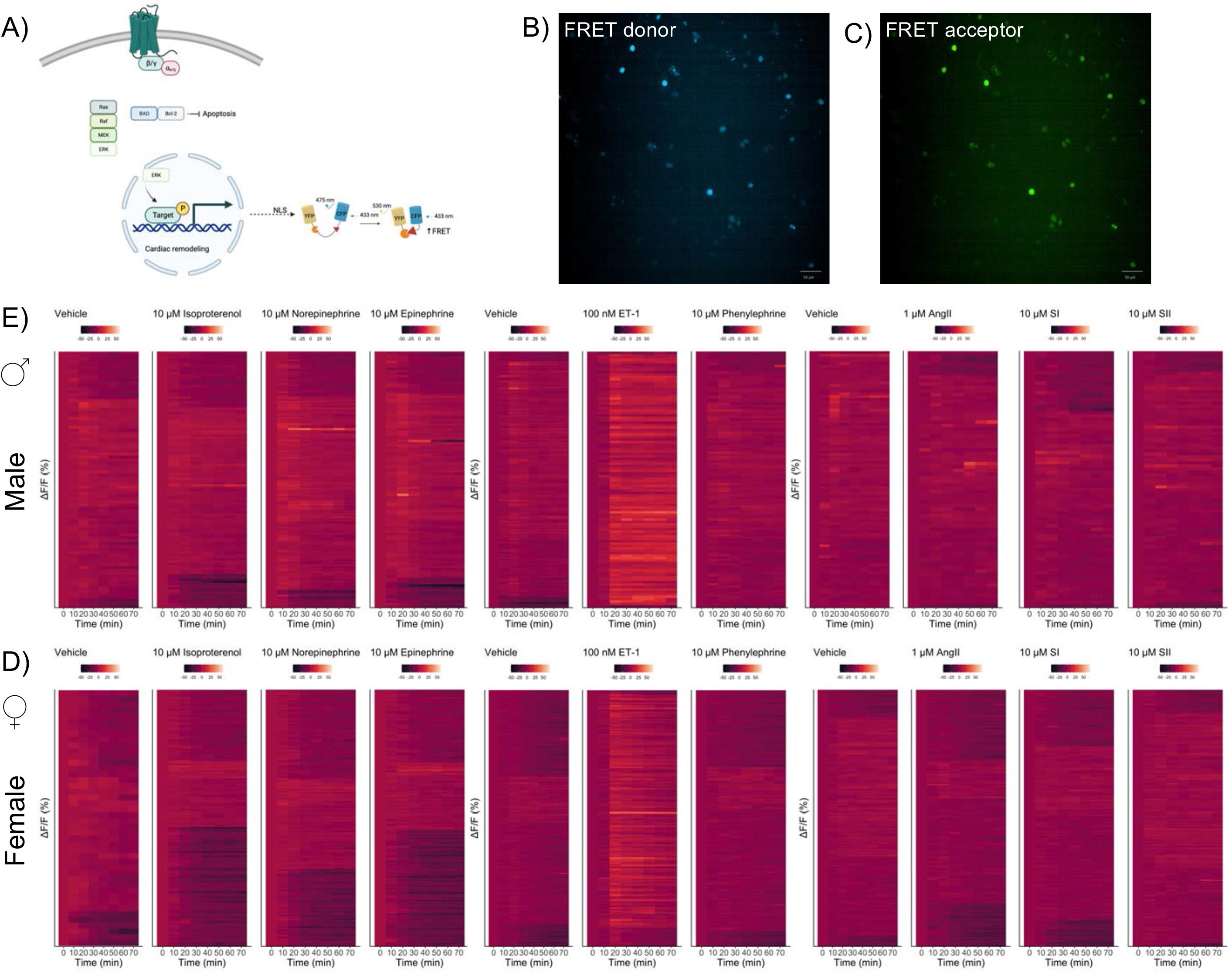
Biosensors to illuminate how biological sex influences cellular signalling. **(A) Schematic illustration of genetically encoded biosensor to interrogate extracellular signal-regulated protein kinase ERK_1/2_ activity.** ERK functions downstream the activation of several receptors and growth factor receptors and regulates cardiomyocyte survival and remodelling. **(B) Fluorescent microscopy demonstrating the nuclear expression of FRET donor and (C) acceptor fluorophores. (D) Representative heatmap representing single cell readouts of FRET-based biosensor EKAREV-NLS in (D) male and (E) female iPSC-CMs, both diagnosed with dilated cardiomyopathy.** The signalosome of two patients, female and male were compared post stimulation by a panel of agonists that target adrenergic, angiotensin and endothelin receptor systems. Adrenergic and angiotensin receptors represent critical pharmacological targets in DCM. Images were collected pre- and post-drug stimulation. Data was collected for 70 minutes post-drug addition. Nuclei from the male patient demonstrated a more pronounced ERK_1/2_ activation profile compared to those of the female patient assayed. While nuclei derived from the female patient exhibited a robust decrease in ERK_1/2_ activity in response to adrenergic drugs.

Overall, our biosensor platform is sensitive enough to detect signalling differences based on biological sex. As HF and DCM progression is known to be sex-specific, our iPSC-biosensor screening platform can help improve cardiac care for females which currently have worse outcomes as majority of studies focus on males (88). Besides biological sex, as our DCM patient cohort includes patients of various ethnicities, this initiative will also be able to shed light on how DCM manifests outside the typical ‘textbook white male’.

## Concluding remarks

The combination of iPSC technology with biosensors provides a unique vantage point geared towards understanding *all* forms of dilated cardiomyopathy and HF. The data created while assessing how control and DCM patient derived iPSC-CMs behave at the cellular, sub-cellular and transcriptomic levels when stimulated with various stressors will create unprecedented knowledge of the molecular underpinnings of various forms of DCM including idiopathic, chemotherapy-induced and inherited. Translating differences in signal transduction or calcium handling may contribute to the identification of novel mechanisms that may be actionable through pharmacological modulation. Likewise, these differences might be distinct and dependent on the specific patient backgrounds and DCM etiology providing mechanistic information on disease and potential disease specific treatment options. The stratification of patients into distinct clusters may lead to the development of treatment strategies that are tailored to patient profiles as opposed to the non-evidence-based prescribing that is currently being performed in the clinic (**Fig. 7**) (89,90). Besides, working with patient-derived iPSC-CMs offers the capacity to discern how biological sex alters dilated cardiomyopathic phenotypes as iPSCs generated from male and female subjects are easily and equally accessible.

**Figure 7.**
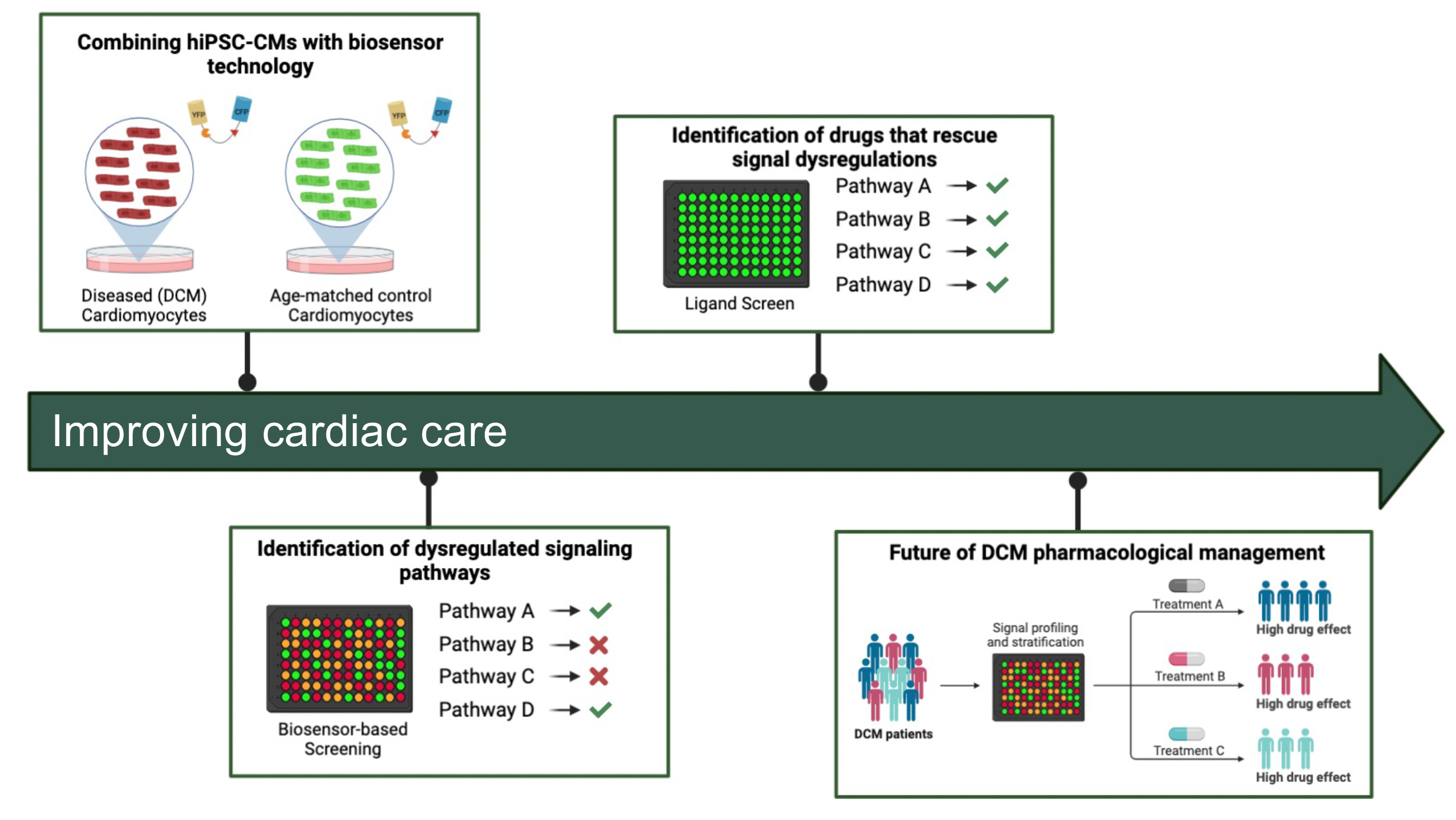
Summary illustration of how biosensor technologies can assist with the improvement of cardiac care. The combination of biosensor technology with iPSC-based models of the cardiomyocyte, either in 2D monolayer or in 3D organoid models as an example of future disease modeling experiments using iPSC-CMs derived from control and patient subjects. The use of genetically encoded biosensors will not only allow for the identification of dysregulated disease-causing pathways but can also facilitate the identification of novel pharmacotherapies to re-establish normal signalling.

Currently, the immaturity of iPSC-CMs poses certain limitations when attempting to extrapolate data for disease modelling. Yet, several publications have already used non-matured iPSC-CMs derived from patients suffering from DCM and these accounts have shown that the iPSC-CMs derived from these patients recapitulate the disease as observed in the clinical setting (**Table 1**). Our analytical pipeline has the integrated capacity to examine maturity-related concerns as morphological features of cardiomyocytes are distinct depending on the maturation stage; neonatal cardiomyocytes are more rounded while adult cardiomyocytes are bi-nucleated and elongated. Our approach, when combined with whole-cell or cytosolic targeted sensors allows to delineate whether distinct signalling profiles are detected in cardiomyocytes of different morphologies and thus developmental stages or maturity. Now, with patient access and the advent of iPSC technology, we can begin to address questions related to DCM progression at molecular level in genetic or idiopathic patient contexts. We believe that the deep phenotyping of these patients using biosensor-based strategies will provide novel insight on disease mechanism that may inform therapeutic interventions.

## Acknowledgements

This work was supported by the Courtois Foundation and grants from the Canadian Institutes of Health Research (CIHR). K.B was supported by a CIHR fellowship and McGill Faculty of Medicine Internal studentship. I.D was supported by a CIHR fellowship and CODE LiFE Foundation Award. C.H was supported by a McGill internal studentship award. J.J.T was supported by fellowships from the CIHR and the McGill Healthy Brains for Healthy Lives initiative. K.A was supported by a Mitacs Accelerate Post-Doctoral Award. A.J was supported by a CIHR master’s award. J.Z. was supported by a Canadian Graduate Student Master’s Award. T.E.H. is the holder of the Canadian Pacific Chair in Biotechnology. The authors would like to thank Dr. Michiyuki Matsuda (Kyoto University) for providing them with the EKAR-EV-NLS biosensor construct and Dr. Jin Zhang for providing the ExRai-AKAR2-NLS biosensor. The authors thank the McGill Pharmacology and Therapeutics Imaging and Molecular Biology platform as well as Dr. Nicolas Audet for microscopy and Columbus support. The authors also thank the McGill Advanced BioImaging facility for training and assistance with microscopy and image analysis. The authors thank members of the Cecere lab for the generation of the iPSC lines. Lastly, the authors thank members of the Hébert laboratory for feedback and guidance throughout the development of the project and for critical reading of the manuscript.

---

**Supplementary Figure 1.**
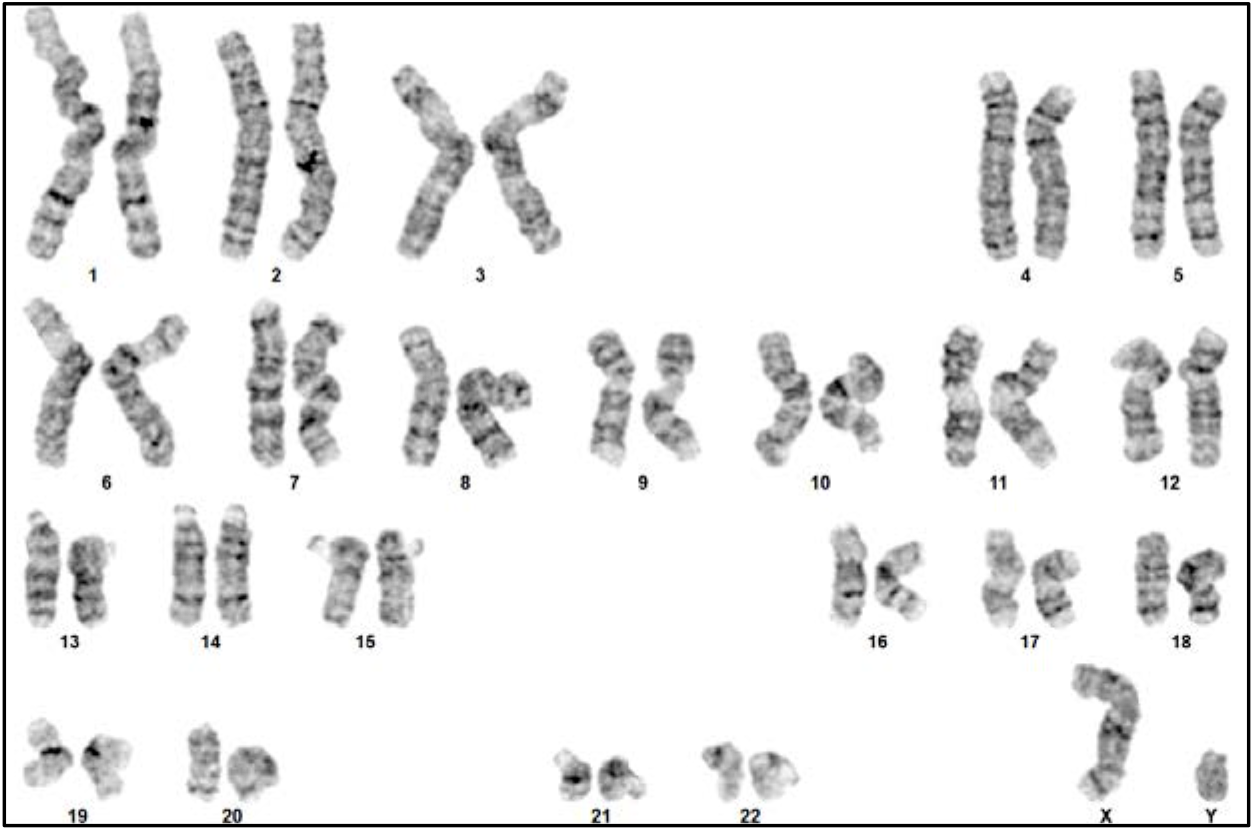
Secondary validation of the induced pluripotent stem cell lines generated. G-band karyotyping analysis of a representative male iPSC cell line. Analysis depicts a normal diploid male karyotype with 23 intact chromosomes. G-banding was performed as means to validate the chromosomal integrity of each iPSC line.

**Supplementary Table 1.** Patient population.

| **Characteristic** | **Patient** | **Control** |
| --- | --- | --- |
| Number of participants (n) | 209 | 19 |
| Age (Mean (SD)) | 54.21 (14.88) | 46.74 (17.94) |
| Male Sex (%) | 143 (69.4) | 7 (38.9) |
| Number of risk factors (Mean (SD)) | 1.8 (1.31) | 1.18 (1.07) |
| Family history (%) | 92 (48.9) | 9 (52.9) |
| Ever smoked (5) | 96 (50.8) | 7 (41.2) |
| High Cholesterol (%) | 49 (25.3) | 2 (11.8) |
| Hypertension (%) | 67 (34.5) | 2 (11.8) |
| Diabetes (%) | 45 (23.2) | 0 (0) |

**Supplementary Table 2.** Summary of heart failure medications prescribed to patient and control subjects

| **Category** | **Number (%) in**<br>**Patients**<br>**(without cardiac transplant)** | **Number (%) in**<br>**Patients**<br>**(with cardiac transplant)** | **Number (%) of**<br>**Controls** |
| --- | --- | --- | --- |
| β-blocker | 166 (88) | 5(25) | 0 (0) |
| Diuretic | 99 (52) | 5(25) | 0 (0) |
| Entresto | 90 (48) | 2(10) | 0 (0) |
| ACE inhibitor | 55 (29) | 5(25) | 0 (0) |
| Angiotensin II receptor Blocker | 20 (11) | 4(20) | 2 (11) |

